# Large cities and the loss of green areas exclude migrant birds: a global analysis

**DOI:** 10.1101/709725

**Authors:** Lucas M. Leveau

## Abstract

Several studies around the world have shown that the proportion of migrant species in bird communities increases toward the poles as a result of greater climatic seasonality and a considerable annual variation of resources. In this context, urban areas may impose a barrier to bird migrants given their buffered seasonality of resources and human disturbance. The aim of this study is to analyze the global pattern of migrant species proportion in urban green areas, considering the effects of climatic seasonality as well as the effects of urbanization. Data of bird communities in urban green areas were gathered through a search of scientific articles, book chapters, and thesis. Datasets that included a list of observed species, the numbers of parks surveyed and other methodological characteristics were considered for the analysis. Then, generalized linear models were used to relate the proportion of migratory species in each dataset to environmental and methodological variables that controlled for different sampling effort among studies. A total of 32 cities from four continents were analyzed. As expected, the migrant proportion increased with the annual range of temperature and precipitation and was higher in the Northern Hemisphere. However, the proportion of migrants decreased with the population size of cities but increased in those datasets with the highest maximum size of green areas surveyed. Although the global pattern of migrant proportion in urban green areas follows a similar pattern than those found in natural areas, the results obtained suggest that urbanization have a negative impact on this global pattern by reducing the proportion of migrant species in large cities. Moreover, green area loss in cities may impact negatively the proportion of migrant species.

## INTRODUCTION

Bird migration is a phenomenon that has attracted considerable attention among ecologists and the general public (Newton 2010). However, its global macrogeographic analysis has been scarcely analyzed (Somveille et al. 2013). Results obtained mainly in the Northern Hemisphere have shown that migrant proportion increases with latitude as a response to seasonal fluctuations of environmental conditions and resources (MacArthur 1959, Herrera 1978, Hulrbert and Haskell 2003, Somveille et al. 2013). Sites near the Poles have a greater seasonal fluctuation of resources that promotes a surplus during spring and summer which is exploited by migrant birds (Herrera 1978, Hurlbert and Haskell 2003). Moreover, findings have shown that bird migration is mainly a Northern hemisphere phenomenon as a result of greater landmass cover at higher latitudes in comparison with the Southern hemisphere (Faaborg et al. 2010, Somveille et al. 2013).

To date, global analyses were based on digital maps of bird species distribution (Somveille et al. 2013, 2019). However, finer-scale data of bird species presence could give more insights about the environmental drivers of bird migration, allowing analyzing possible effects of land-use change on migrant birds (Zuckerberg et al. 2016, La Sorte et al. 2017).

Populations of migratory bird species have been declining, at least for the Northern Hemisphere where long term data is available (Robbins et al. 1988, Bailie and Peach 1992, Berthold et al. 1998). Causes of migrant declines have turned out to be related with land-use change in breeding and wintering grounds during 1970-1990 to climate and land-use change in the last decades (Møller et al. 2008, Newton 2008, Cox 2010, Morrison et al. 2013). The advance of spring greening promoted by climate warming in the Northern Hemisphere is associated with a mismatch between the peak of food availability and the timing of breeding, which is postulated as the main driver of migrant population declines (Both et al. 2006, Møller et al. 2008, Cox 2010).

Since 2008 more than half of the human population lives in cities (Handwerk 2008) and there is an increasing trend of population growth in the next years (United Nations 2018). This population growth is related to urban expansion over natural and agricultural areas and the increase of building cover and loss of green areas within cities (Dallimer et al. 2011; Vergnes et al. 2014). The loss of green areas can affect negatively to migrant species richness and abundance (Mason et al. 2007, Palmer et al. 2008, Pennington et al. 2008, Zhou and Chou 2012, Leveau 2013, Kang et al. 2015). The level of urbanization surrounding green areas also can impact migrant individual’s proportion and richness (Friesen et al. 1995, Hennings and Edge 2003, Dunford and Freemark 2004, Rodewald and Matthews 2005, Mason et al. 2007, Pennington et al. 2008). Moreover, migrant species seem more sensitive than resident species to green area size and human disturbance (Burger and Gochfeld 1991, Park and Lee 2000, Zhou and Chu 2012). Given that urban areas tend to have a lower seasonal fluctuation of primary productivity induced by the urban heat island phenomenon, the food resources surplus during spring-summer may be lower than non-urban areas, affecting the presence of migratory species in cities (Leveau et al. 2018, Leveau 2018). In addition, the urban heat island phenomenon can exacerbate the mismatch between the peak of food availability and the timing of breeding of migrant species.

The aims of this study are: 1) to analyze the relationship between migrant species proportion in urban green areas of cities located worldwide and climatic variation and hemisphere; 2) to relate migrant proportion with the size of green areas; and 3) to analyze the relationship between migrant proportion and urbanization, assessed as city size and city age. I collected studies of bird communities in urban green areas and controlled for bird sampling methodology and sampling effort. I expected that migrant proportion will be the highest in cities located near the poles, where annual climatic fluctuation is prominent, and highest in the Northern Hemisphere. Moreover, an increased migrant proportion is expected for studies that included the largest green areas sizes, and for studies conducted in the oldest cities because these are expected to have more habitat complexity in their green areas. Finally, I expected depletion in migrant proportion in the largest cities, because these have an increase of car and pedestrian traffic, pollution, more isolation to non-urban areas and a lower seasonality of resources, which potentially affect negatively to migrant species.

## METHODS

### Data collection

During September 2015 several databases, such as Scopus and Google scholar, were used to find published articles and chapters with the following key-words: [avian OR bird*] AND [green OR park OR cemeter* OR remnant* OR golf] AND urban, in English as well as in Spanish. Moreover, I searched for additional articles in reviews of urban birds (Fernández-Juricic and Jokimäki 2001; Chace and Walsh 2006; Garden et al. 2006; Ortega-Alvarez and MacGregor-Fors 2011).

Urban green areas were defined as patches dominated by vegetation and surrounded by a matrix of built-up land uses (Matthews et al. 2015). This definition included urban parks, remnants of native vegetation, golf courses, and cemeteries. This analysis was focused only on breeding bird communities, i.e. surveyed during spring-summer seasons, because this period of the year is the most analyzed among studies (Leveau et al. 2019). Only papers that contained the following data were considered: 1) a list of species found; 2) the number of surveys made to each green area; 3) the number of green areas surveyed; 4) the maximum size of the green areas surveyed; and 5) the method of bird survey. In the case to obtain several studies from the same city, only the study with more green areas surveyed was considered. The list of species was employed to calculate the proportion of latitudinal migrant species in each study. If the information of the migratory status of each species was not provided in the paper, status data were obtained from the bibliography (Yamashina 1961, Peterson et al 1973, Wild Bird Society of Japan 1982, National Geographic Society 1999; del Hoyo et al. 1994-2011). Variables 2) and 3) were included to control for sampling effort, given that more visits to each green area and more green areas surveyed may include more migrant species than less visits and green areas surveyed. The maximum size of the green areas surveyed in each study was included because of species-area relationships, in which at larger area size it is expected a greater number of migrant species. The method of bird survey was classified in point counts, transects count or other methods, which included the mapping method or a combination of transect and point counts. Other variables, such as city population size, the altitude of the city and foundation were obtained from papers; otherwise, data was gathered from Wikipedia. Population data from the nearest year to the bird surveys was considered.

Studies which only focused on a certain bird assemblage (e.g., forest specialists or migrant birds) or that excluded some species, such as the rock dove *(Columba livia),* were excluded from the analysis. However, studies that excluded waterfowl were included given that in general, these birds are in low numbers in green areas (Murgui 2010). Some studies only considered the breeding birds, and given that in an area there could be non-breeding migrant species (e.g. the Barn swallow *[Hirundo rustica*] in South America during the austral spring-summer), these studies could affect the results obtained. Therefore, this category of datasets also was included as a control variable in the statistical model.

The following climatic variables for each city were obtained from Wikipedia: 1) minimum annual temperature, TMIN (°C); 2) maximum annual temperature, TMAX; 3) minimum annual precipitation, PMIN (mm); 4) and maximum annual precipitation, PMAX. Then I calculated the annual range of temperature (TRANGE), the annual range of precipitation (PRANGE), the annual relative change of temperature (TRELATIVE = TRANGE/TMAX), and the annual relative change of precipitation (PRELATIVE= PRANGE/PMAX).

### Statistical analysis

Given that the eight climatic variables were highly correlated, a principal component analysis (PCA) was made to summarize them with latitude in a few axes of climatic variation among cities. The function princomp of R (The R development team 2018) was used. The axes with eigenvalues greater than 1 were considered as main climatic factors (Kaiser 1960).

The climatic axes, the methodological and the environmental characteristics of cities were related to the proportion of migrant species by a Generalized Linear model with a binomial distribution of errors using the glm function in R. The proportion of migrant species was obtained with the function cbind using the total bird richness and migrant richness. The best model was obtained with the function stepAIC of the MASS package (Ripley 2018). The correlation between continuous explanatory variables was not important (r < 0.70). Plots of the regression models were constructed with the visreg package (Breheny and Burchett, 2013).

## RESULTS

Of 119 articles screened that analyzed bird communities in urban green areas, relevant information was found for 32 cities located in Europe (37%), North America (31%), South America (16%) and Asia (16%) (Figure 1). Migrant proportion varied from 0 to 0.77 (mean = 0.33).

**Figure 1.**
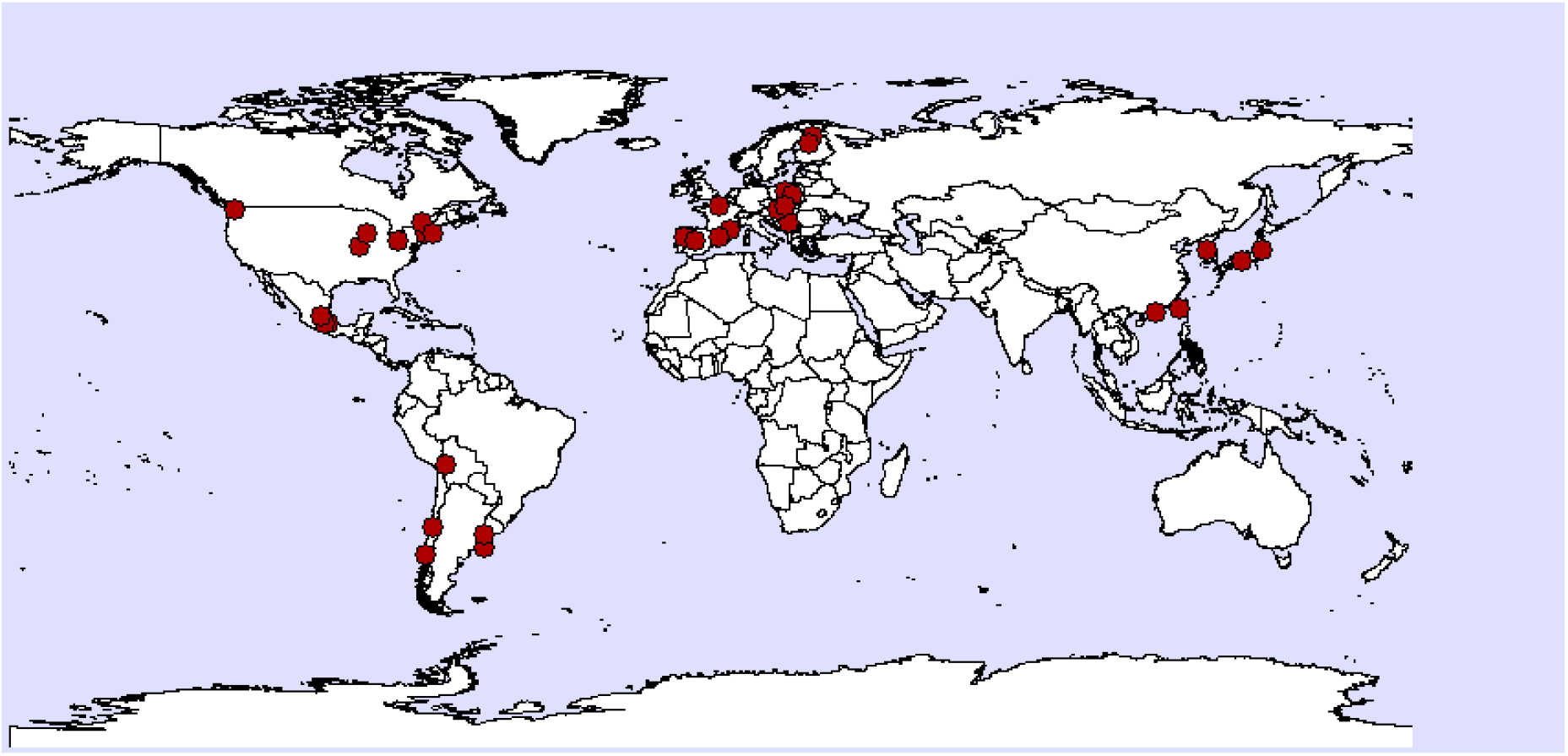
Location of the cities analyzed in this study.

PCA revealed three axes of climatic variation that accounted for 90 % of the variance (Table 1). The first axis was positively related to the annual change and the relative change of precipitation, and negatively related to the annual change and the relative change of temperature. The second axis was positively related to the maximum annual temperature and the annual minimum precipitation. Finally, the third axis was negatively related to the maximum annual precipitation and annual precipitation range. GLMs revealed that the hemisphere, climatic variables, city population and the number of green areas surveyed in each city were related to the migrant proportion (Table 2). Temperature annual range and relative change, which increased at higher latitudes, were positively related to the proportion of migratory species (Figure 2). The annual change of precipitation was positively related to the proportion of migrants (Figure 2). Moreover, the proportion of migrant species was higher in the Northern hemisphere (Figure 3). A greater sampling effort, indicated by the number of green areas visited in each city, also was related to an elevated proportion of migratory species (Figure 4).

**Figure 2.**
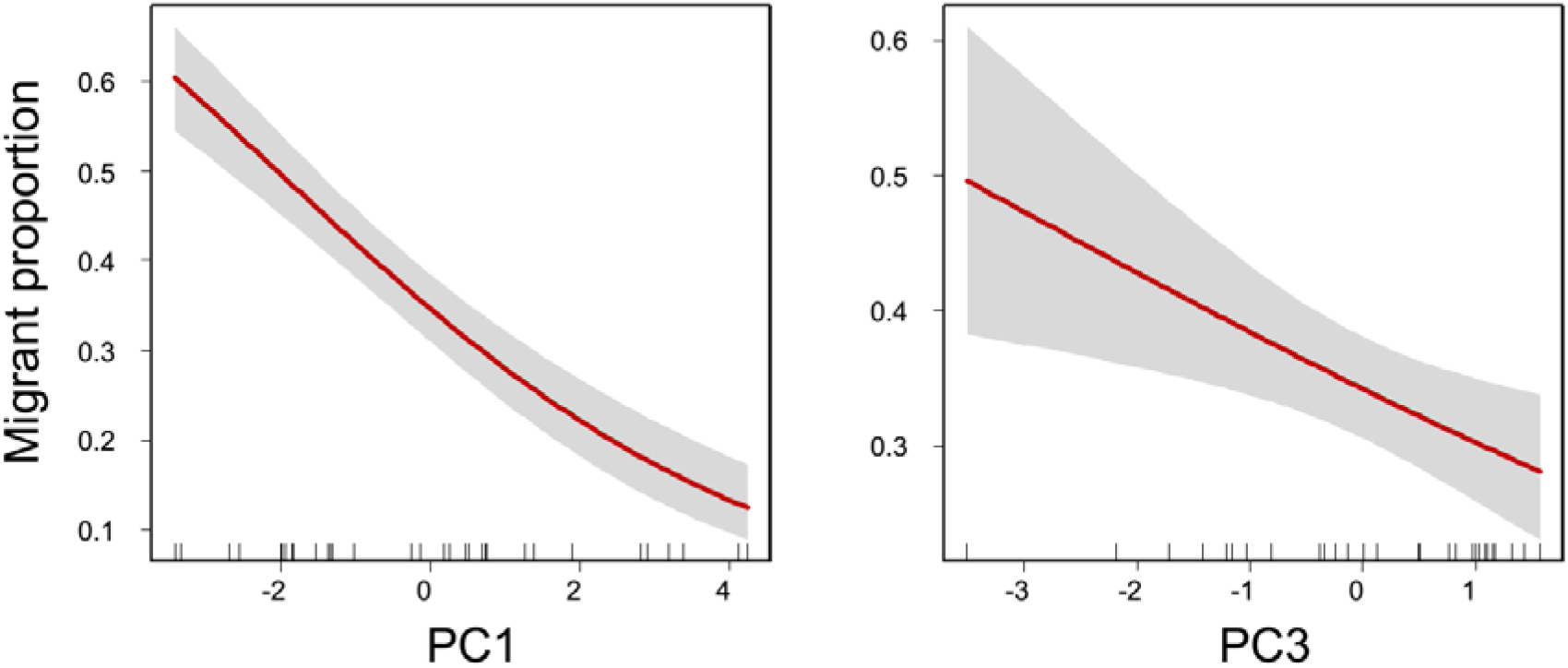
Relationship between climatic axes and the proportion of migrant species in urban green areas. The red line represents the fitted curve and the grey areas are the confidence intervals at 95%.

**Figure 3.**
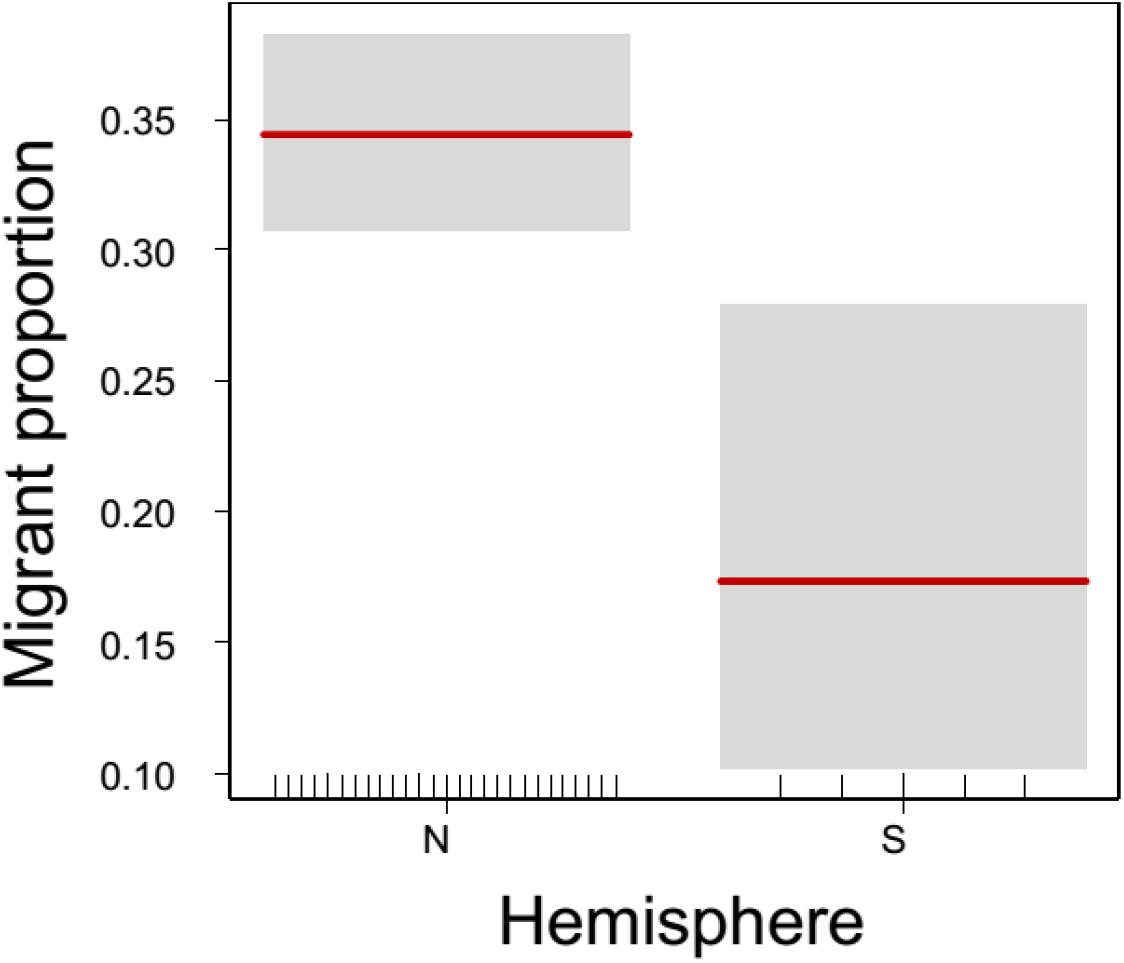
Migrant proportion in the Northern and Southern hemispheres. The red line represents the mean and the grey areas are the confidence intervals at 95%.

**Figure 4.**
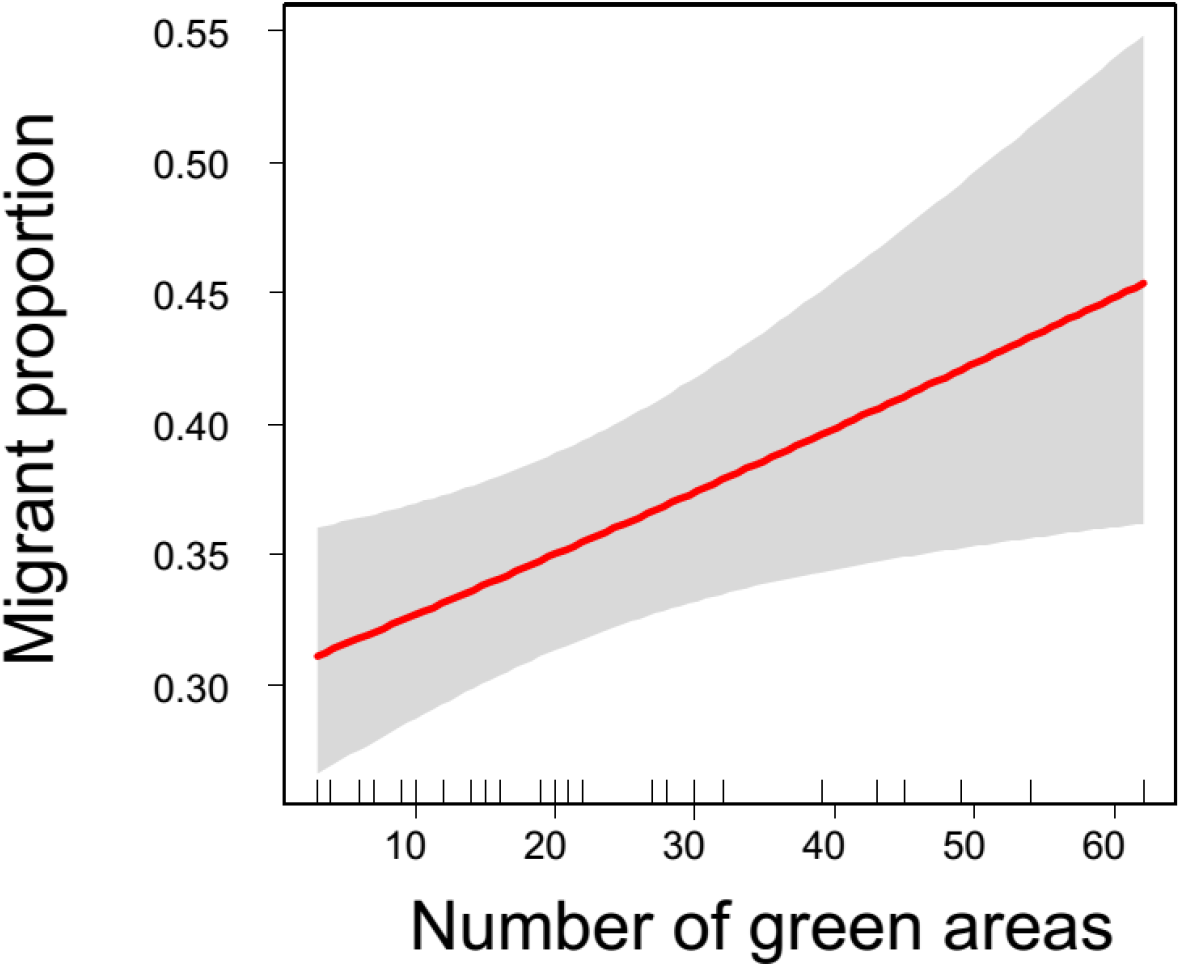
Relationship between migrant proportion and the number of green areas in each dataset. The red line represents the fitted curve and the grey areas are the confidence intervals at 95%.

**Table 1.** Principal component analysis of climatic variables for the 32 cities analyzed. TMIN: minimum annual temperature (°C); TMAX: maximum annual temperature; PMIN: minimum annual precipitation (mm); PMAX: maximum annual precipitation; TRANGE: annual range of temperature; PRANGE: the annual range of precipitation; TRELATIVE: the annual relative change of temperature; and PRELATIVE: the annual relative change of precipitation.

| Variables | PC1 | PC2 | PC3 |
| --- | --- | --- | --- |
| TMIN | 0.69 | 0.59 | 0.36 |
| TMAX | -0.06 | 0.93 | 0.08 |
| PMIN | -0.61 | 0.62 | -0.10 |
| PMAX | 0.65 | 0.28 | -0.69 |
| PRANGE | -0.77 | 0.46 | -0.29 |
| TRANGE | 0.74 | 0.15 | -0.65 |
| TRELATIVE | -0.86 | -0.26 | -0.41 |
| PRELATIVE | 0.83 | -0.38 | -0.13 |
| LATITUD | -0.84 | -0.22 | -0.18 |
| Eigenvalues | 4.56 | 2.18 | 1.35 |
| % Variance | 50.63 | 24.23 | 15.03 |

**Table 2.** Best model explaining the proportion of migrant species in green areas of cities worldwide.

|  | Estimate | SE | z value | P |
| --- | --- | --- | --- | --- |
| Intercept | -0.809 | 0.125 | -6.486 | 0.000 |
| PC1 (climate) | -0.310 | 0.036 | -8.697 | 0.000 |
| PC3 (climate) | -0.182 | 0.064 | -2.825 | 0.005 |
| Population (millions) | -0.095 | 0.021 | -4.472 | 0.000 |
| Number of green areas | 0.010 | 0.004 | 2.435 | 0.015 |
| Maximum size of green areas (Ha) | 0.002 | 0.001 | 2.341 | 0.019 |
| Hemisphere (South) | -0.919 | 0.330 | -2.785 | 0.005 |

The increase of city size was related to a decrease of migrant proportion, whereas the maximum size of the green areas surveyed had a positive effect on the migrant proportion (Figure 5).

**Figure 5.**
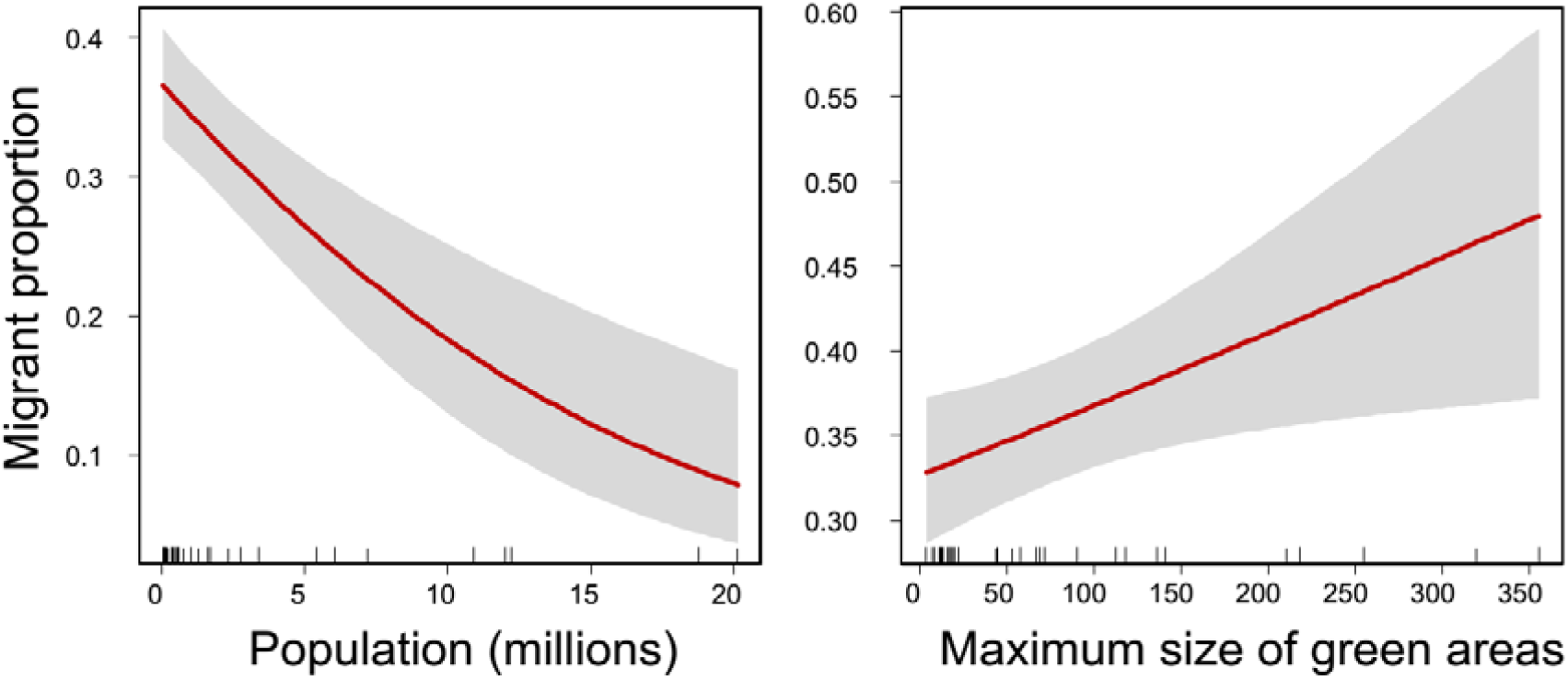
Relationship between migrant proportion, population size of cities and the maximum size of green areas in each dataset. The red line represents the fitted curve and the grey areas are the confidence intervals at 95%.

## DISCUSSION

The results obtained showed that, as expected, the proportion of migrant species in each assemblage increased with latitude and the annual variation of temperature and precipitation. The greatest variation of climatic conditions may be related to annual resource variation and the surplus exploited by migrant species. Furthermore, the proportion of migrant birds was affected by city population size and green area size, suggesting that urbanization negatively impacts the use of migratory species of urban green areas.

As pointed out by several authors (MacArthur 1959, Willson 1976, Herrera 1978, Newton and Dale 1996a,b, Somveille et al. 2013), the migrant proportion increases with latitude and it depends on the magnitude of the absolute change of resource availability between breeding and nonbreeding seasons. Given that most of the migrant species feed on insects, temperature annual changes may be responsible for their fluctuations, promoting seasonal “blooms” exploited partially by the migrant arrival (Herrera 1978, Hurlbert and Haskell 2003). The results obtained in this study strongly agree with previous findings, given that migrant proportion had a tight positive relationship with latitude and annual changes in temperature.

Hemisphere influenced migrant proportion, which increased in the Northern hemisphere. This result agrees with other studies which postulated that a greater landmass in high latitudes of the Northern hemisphere compared to the Southern hemisphere induced the presence of a higher proportion of migrant species (Faaborg et al. 2010, Somveille et al. 2013).

Larger cities had the lowest proportion of migrant species, suggesting that urbanization have a negative impact on the global patterns of bird migration. As cities grow, they are characterized by higher levels of car and pedestrian traffic (Leveau and Leveau 2006), which negatively affect migrant proportion and richness (Burger and Gochfeld 1991, Hennings and Edge 2003, Zhou and Chu 2012). Moreover, more isolation of green areas to non-urban areas and a lower seasonality of resources can decrease migrant bird presence (Leveau and Leveau 2006, Leveau 2018, Leveau et al. 2018). Finally, as cities grow urban green areas can become surrounded by denser urbanization, which may decrease migrant richness and abundance (Friesen et al. 1995, Dunford and Freemark 2004, Pennington et al. 2005, Rodewald and Matthews 2005, Mason et al. 2007). Although the number of bird species in urban green areas increases with area size consistently worldwide (Leveau et al. 2019), several studies made at local scales have shown that migrant species are more sensitive to green area size than resident species (Park and Lee 2000, Zhou and Chu 2012). The results obtained in this study indicated that the maximum size of green areas in each dataset increased migrant proportion, suggesting that green area loss in cities may affect negatively to migrant birds.

Although this study was focused on bird assemblages surveyed during spring-summer, other analyses conducted in eastern North America during an annual cycle found that migratory bird species are positively associated to urban areas during autumn (La Sorte et al. 2017). Therefore, it is necessary a more detailed analysis of the annual use of urban green areas by migrant bird species.

## CONCLUSIONS

The results obtained showed that the proportion of migrant bird species follow a similar global pattern than in natural areas, increasing their relative numbers toward areas with the greatest seasonal variation of climate and being more predominant in the Northern hemisphere. However, urbanization seems to have a negative impact on this global pattern by reducing the proportion of migrant species in large cities. Moreover, green area size may be an important aspect that promotes the entry of migrant species in cities. Given that migratory species are a substantial part of the seasonal dynamics of bird communities, the loss of migrant proportion in large cities supports the idea that urbanization causes a temporal homogenization of bird communities (Leveau 2018). It is necessary to conduct multiscale studies to get more insights about urban green area use by migratory species, producing the knowledge necessary to foster management strategies that mitigate the negative impacts of cities on migrant species.

## References

Baillie, S. R., & Peach, W. J. 1992. Population limitation in Palaearctic-African migrant passerines. Ibis 134: 120–132.

Berthold, P., Fiedler, W., Schlenker, R., & Querner, U. 1998. 25-year study of the population development of Central European songbirds: a general decline, most evident in long-distance migrants. Naturwissenschaften 85: 350–353.

Both, C., Bouwhuis, S., Lessells, C. M., & Visser, M. E. 2006. Climate change and population declines in a long-distance migratory bird. Nature 441: 81.

Breheny, P., & Burchett, W. 2013. Visualizing regression models using visreg. http://myweb.uiowa.edu/pbreheny/publications/visreg.pdf

Burger, J., & Gochfeld, M. 1991. Human distance and birds: tolerance and response distances of resident and migrant species in India. Environ Conserv 18: 158–165.

Chace, J. F., & Walsh, J. J. 2006. Urban effects on native avifauna: a review. Landsc Urban Plann 74: 46–69.

Cox, G. W. (2010). Bird migration and global change. Island Press.

Dallimer, M., Tang, Z., Bibby, P. R., Brindley, P., Gaston, K. J., & Davies, Z. G. 2011. Temporal changes in greenspace in a highly urbanized region. Biol Lett 7: 763–766.

Del Hoyo J, Elliott A, Christie D (1994-2011) Handbook of the birds of the world. Lynx editions, Barcelona

Faaborg, J., Holmes, R. T., Anders, A. D., Bildstein, K. L., Dugger, K. M., Gauthreaux, S. A., … & Latta, S. C. (2010). Conserving migratory land birds in the New World: Do we know enough? Ecol Appl 20: 398–418.

Fernandez-Juricic, E., & Jokimäki, J. 2001. A habitat island approach to conserving birds in urban landscapes: case studies from southern and northern Europe. Biodivers Conserv 10: 2023–2043.

Friesen, L. E., Eagles, P. F., & Mackay, R. J. 1995. Effects of residential development on forest-dwelling Neotropical migrant songbirds. Conserv Biol 9: 1408–1414.

Garden, J., Mcalpine, C., Peterson, A. N. N., Jones, D., & Possingham, H. 2006. Review of the ecology of Australian urban fauna: a focus on spatially explicit processes. Aust Ecol 31: 126–148.

Handwerk, D. 2008. Half of humanity will live in cities by year’s end. http://news.nationalgeographic.com/news/2008/03/080313-cities.html

Hennings, L. A., & Edge, W. D. 2003. Riparian bird community structure in Portland, Oregon: habitat, urbanization, and spatial scale patterns. Condor, 105: 288–302.

Herrera, C. M. 1978. On the breeding distribution pattern of European migrant birds: MacArthur’s theme reexamined. Auk 95: 496–509.

Hurlbert, A. H., & Haskell, J. P. 2002. The effect of energy and seasonality on avian species richness and community composition. Am Nat 161: 83–97.

Kaiser, H. F, 1960. The application of electronic computers to factor analysis. Educ Psychol Meas 20:141–151

Kang, W., Minor, E. S., Park, C. R., & Lee, D. 2015. Effects of habitat structure, human disturbance, and habitat connectivity on urban forest bird communities. Urban Ecosyst 18: 857–870.

La Sorte, F. A., Fink, D., Buler, J. J., Farnsworth, A., & Cabrera-Cruz, S. A. 2017. Seasonal associations with urban light pollution for nocturnally migrating bird populations. Global Change Biol 23: 4609–4619.

Leveau, L. M. 2013. Relaciones aves–habitat en el sector suburbano de Mar del Plata, Argentina. Ornitol Neotrop 24: 201–212.

Leveau, L. M. 2018. Urbanization, environmental stabilization and temporal persistence of bird species: a view from Latin America. PeerJ 6: e6056.

Leveau, L. M., Isla, F. I., & Bellocq, M. I. 2018. Predicting the seasonal dynamics of bird communities along an urban-rural gradient using NDVI. Landsc Urban Plann 177: 103–113.

Leveau, L. M., A. Ruggiero, T. Matthews, Bellocq M. I. 2019. A global consistent positive effect of green area size on bird richness. Avian research.

Mason, J., Moorman, C., Hess, G., & Sinclair, K. 2007. Designing suburban greenways to provide habitat for forest-breeding birds. Landsc Urban Plann 80: 153–164.

Møller, A. P., Rubolini, D., & Lehikoinen, E. 2008. Populations of migratory bird species that did not show a phenological response to climate change are declining. Proc Nat Acad Sci 105: 16195–16200.

Morrison, C. A., Robinson, R. A., Clark, J. A., Risely, K., & Gill, J. A. (2013). Recent population declines in Afro-Palaearctic migratory birds: the influence of breeding and non-breeding seasons. Divers Distrib 19: 1051–1058.

National Geographic Society (US). 1999. Field guide to the birds of North America. National Geographic, New York

Newton I (2010) The migration ecology of birds. Elsevier, London.

Newton, I., & Dale, L. 1996a. Relationship between migration and latitude among west European birds. J Anim Ecol: 137–146.

Newton, I., & Dale, L. C. 1996b. Bird migration at different latitudes in eastern North America. Auk 113: 626–635.

Ortega-Álvarez, R., & MacGregor-Fors, I. 2011. Dusting-off the file: A review of knowledge on urban ornithology in Latin America. Landsc Urban Plann 101: 1–10.

Palmer, G. C., Fitzsimons, J. A., Antos, M. J., & White, J. G. 2008. Determinants of native avian richness in suburban remnant vegetation: implications for conservation planning. Biol Conserv 141: 2329–2341.

Park, C. R., & Lee, W. S. 2000. Relationship between species composition and area in breeding birds of urban woods in Seoul, Korea. Landsc Urban Plann 51: 29–36.

Pennington, D. N., Hansel, J., & Blair, R. B. 2008. The conservation value of urban riparian areas for landbirds during spring migration: land cover, scale, and vegetation effects. Biol Conserv 141: 1235–1248.

Peterson, R., Mountfort, G., Hollom, P. A. D., & Díaz, G. 1973. Guía de campo de las aves de España y demás países de Europa (No. C/598.294 P4).

Ripley, B. 2011. MASS: support functions and datasets for Venables and Ripley’s MASS. R package version, 7–3.

Robbins, C. S., Sauer, J. R., Greenberg, R. S., & Droege, S. 1989. Population declines in North American birds that migrate to the Neotropics. Proc Nat Acad Sci 86: 7658–7662.

Somveille, M., Manica, A., Butchart, S. H., & Rodrigues, A. S. 2013. Mapping global diversity patterns for migratory birds. PloS one 8: e70907.

Somveille, M., Manica, A., & Rodrigues, A. S. (2019). Where the wild birds go: explaining the differences in migratory destinations across terrestrial bird species. Ecography, 42(2), 225–236.

United Nations. (2018). World urbanization prospects: The 2018 revision. Retrieved from https://esa.un.org/unpd/wup/Publications/Files/WUP2018-KeyFacts.pdf

Vergnes, A., Pellissier, V., Lemperiere, G., Rollard, C., & Clergeau, P. 2014. Urban densification causes the decline of ground-dwelling arthropods. Biodivers and Conserv 23: 1859–1877.

Wild Bird Society of Japan (1982) A field guide to the birds of Japan. Kodansha International Limited, Tokyo.

Willson, M. F. 1976. The breeding distribution of North American migrant birds: a critique of MacArthur (1959). Wilson Bull 88: 582–587.

Yamashina Y (1961) Birds in Japan: a field guide. Tokyo news Limited, Tokyo.

Zhou, D., & Chu, L. M. 2012. How would size, age, human disturbance, and vegetation structure affect bird communities of urban parks in different seasons? J Ornithol 153: 1101–1112.

Zuckerberg, B., Fink, D., La Sorte, F. A., Hochachka, W. M., & Kelling, S. 2016. Novel seasonal land cover associations for eastern North American forest birds identified through dynamic species distribution modelling. Divers Distrib 22: 717–730.

